# Two distinct feedback codes in V1 for ‘real’ and ‘imaginary’ internal experiences

**DOI:** 10.1101/664870

**Authors:** Johanna Bergmann, Andrew T. Morgan, Lars Muckli

## Abstract

Visual illusions and visual imagery are conscious sensory events that lack a corresponding physical input. But while everyday mental imagery feels distinct from incoming stimulus input, visual illusions, like hallucinations, are under limited volitional control and appear indistinguishable from physical reality. Illusions are thought to arise from lower-level processes within sensory cortices. In contrast, imagery involves a wide network of brain areas that recruit early visual cortices for the sensory representation of the imagined stimulus. Here, we combine laminar fMRI brain imaging with psychophysical methods and multivariate pattern analysis to investigate in human participants how seemingly ‘real’ and imaginary non-physical experiences are processed in primary visual cortex (V1). We find that the content of mental imagery is only decodable in deep layers, whereas illusory content is only decodable at superficial depths. This suggests that feedback to the different layers may serve distinct functions: low-level feedback to superficial layers might be responsible for shaping perception-like experiences, while deep-layer feedback might serve the formation of a more malleable ‘inner’ world, separate from ongoing perception.

## Main text

During visual illusions, we might see an object that is not physically present(*1, 2*), *not* see an object that is physically present(*3*) or perceive an object’s physical properties to be different to how they actually are(*4*). Such illusions are attributed to relatively low-level processes, such as lateral interactions within primary visual cortex(*4, 5*) or feedback from other visual areas(*6*). Higher-order visual areas have larger receptive fields and can thus integrate input from larger portions of the visual field. Based on this larger-scale arrangement, we can make guesses about the global characteristics of individual objects, which are fed back to lower-order areas like V1. This process speeds up and facilitates perception(*7*), but can also lead to striking misperceptions. For example, during the Kanizsa or the neon colour-spreading illusions, we see illusory contours of a (coloured) shape that is physically not there. This experience is thought to arise because higher-order visual areas assume the presence of a shape based on the alignment of other shapes in the visual field(*8, 9*).

Just like hallucinations, illusions typically seem ‘real’, and our volitional control of their experience is limited; they appear embedded within the external environment, indistinguishable from physical reality. By contrast, everyday mental imagery feels clearly distinct and segregated from reality and can influence perception and vice versa(*10, 11*). In fact, we can engage in both simultaneously – we can freely form visual thoughts that guide our actions and decisions, while simultaneously processing perceptual input from our senses.

A widely distributed network of brain areas is involved in imagery. Evidence suggests that high-level frontoparietal regions recruit early visual cortex for the sensory representation of a mental stimulus(*12*). Connectivity to V1 follows several basic rules. For example, (most) feedforward sensory input from the eye arrives in the middle layers, whereas (most) feedback input from other brain areas arrives in the deep and superficial layers(*13, 14*). On this basis, we hypothesized that information on non-physical visual experiences, like imagery and illusions, should be predominant in the layers that receive feedback from other brain areas. We also wanted to explore whether low-level visual illusions, which arise from feedback within visual cortex, show different layer-wise information profiles relative to mental imagery, which involves feedback from a widely-distributed network. Long-distance feedback from distant brain regions travels through deep layers, with some bifurcations also targeting superficial layers. In contrast, short-range feedback axons that connect to nearby regions are located in and target superficial layers(*13, 15, 16*).

Here, we used high-resolution laminar fMRI (0.8mm^3^) in human participants to explore how the content of illusory perception and visual imagery is represented at different depths of V1. During the fMRI measurement, participants maintained fixation while they were presented with five conditions (**Fig. 1A & B** and Methods): three conditions measured activity patterns during mental imagery, perception, and illusory perception; two conditions acted as controls for the illusory perception condition. For the mental imagery task, we instructed participants to imagine a central red or green disc. A faint grey circle on the screen indicated the visual size and location of the stimulus that participants were asked to imagine. Two seconds prior to the imagery phase, we presented participants with a letter for one second to indicate which colour to imagine. The cue appeared at random locations outside of the location of imagery, with its distance to the fixation cross held constant. In the perceptual condition, participants viewed a central red- or green-coloured disc of the same size as the central circle used in the mental imagery condition. During the illusory perception condition, participants viewed the neon colour spreading illusion, in which four pacman-like ring stimuli induce the illusion that a red or green square shape sits between them (**Fig. 1A**). The peripheral pacman-like stimuli were located outside of the central circle used in the imagery condition to indicate the location of imagery. In the first control of the illusory perception condition, we presented participants with an ‘amodal’ version of the stimulus, in which a white contour was placed in the area between the pacman-like rings. This contour breaks or attenuates the illusory experience, or can be perceived as overlaying the (illusory) shape behind it. The second control consisted of a mock version of the illusion, in which the coloured quarter of the rings were rotated outwards, such that no illusory shape arises.

**Figure 1.**
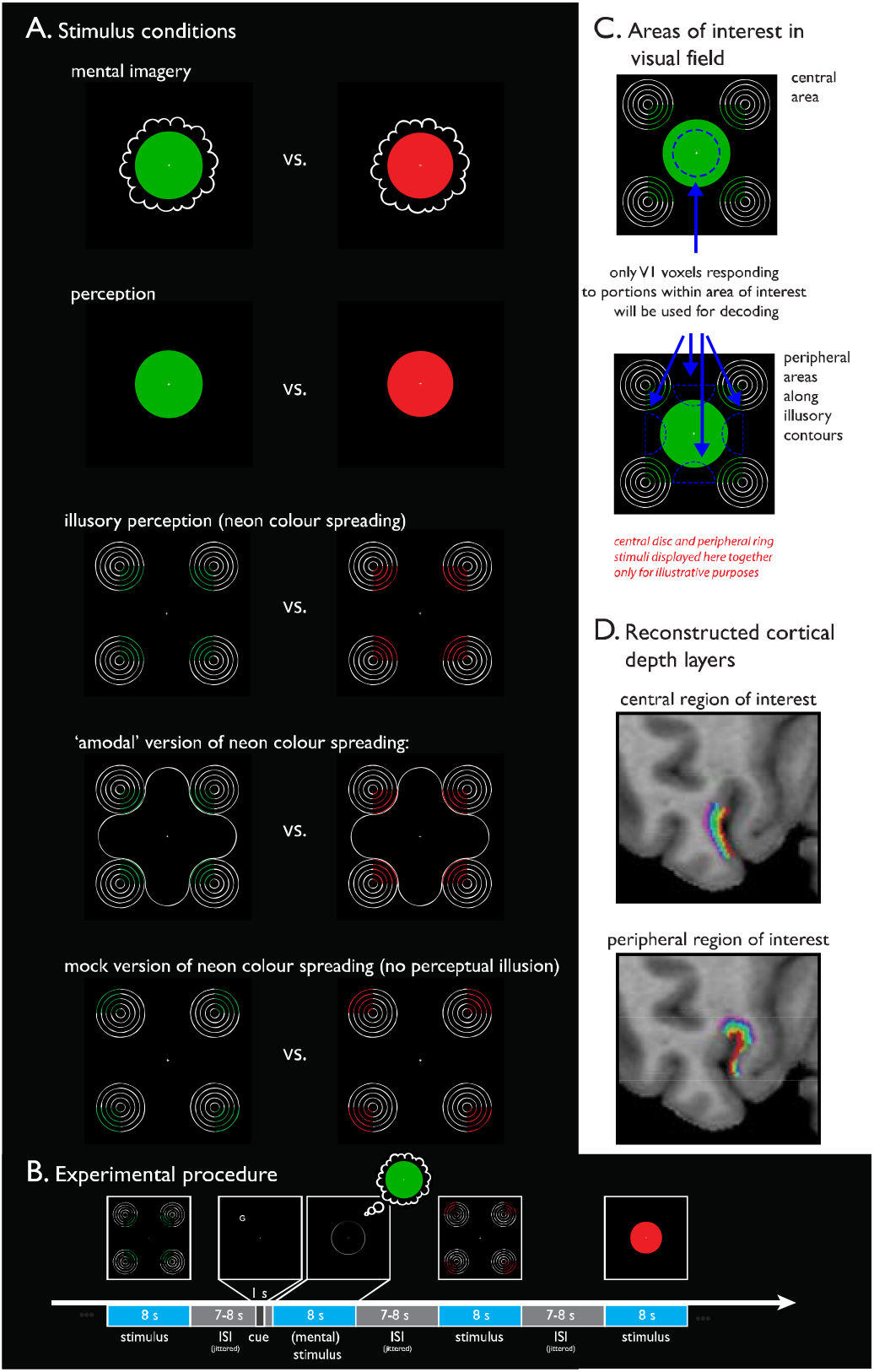
Experimental stimulation and procedure, visual areas of interest and cortical depth layer segmentation. **A.** Five experimental conditions were used: mental imagery, perception, illusory perception and 2 additional control conditions of the illusory perception condition, which comprised ‘amodal’ and mock versions of the stimuli. The stimuli of the five conditions used were in red or green to enable colour to be decoded within each condition. **B.** Each stimulus was presented for 8 seconds in a randomized order. During mental imagery, only a faint grey circle was presented to indicate the location of where participants should imagine the colour. A letter cue preceding the imagery phase indicated which colour to imagine. It was shown at random locations outside of the field of imagery, with the distance to the fixation cross held constant. **C.** We identified two V1 regions of interest (ROI) based on the visual field portions they represented (dashed blue lines). Note that the pac-man like rings of the illusory perception condition and the central colour disc were never presented at the same time and are only displayed here together for illustrative purposes. **C.** In each V1 ROI, the grey matter was segmented into 6 cortical depth layers, which represent coarse approximations of anatomical layers; neighbouring layers may partly share the same voxels.

Individuals vary immensely in their ability to form mental images. To increase the probability of finding imagery-related signals in V1, we focused on those individuals with good imagery abilities. We therefore pre-screened our participants with a behavioural method that determines individual imagery by measuring its impact on subsequent perception(*11*) (Methods and **Supplementary Fig. 1**). Only individuals with imagery scores at or above an *a priori* defined threshold were invited to participate in the fMRI session.

We were particularly interested in cortical processes around two areas in the visual field: (1) the central area around the fixation cross, i.e. the central portion of the area where the participants imagined a coloured disc in the mental imagery task; and (2) four portions in the visual periphery, where the illusory contours of the illusion are located (dashed blue lines in **Fig. 1C**). To identify the V1 portions that process input from these visual field areas, we estimated each voxel’s population-receptive field from visual field mapping stimulation(*17*), and used this approach to define regions of interest (ROI) for further analysis (**Supplementary Fig. 2 & 3**; Methods). We then segmented the identified V1 regions into six cortical depth layers (**Fig. 1D**). For each cortical depth, we used multivariate pattern analysis to determine whether we can detect information about illusory or imagined content. To do so, we trained a support vector machine (SVM) classifier to distinguish the red and green colour of the (illusory/imagined/physical) stimuli in each of the five conditions.

As mental imagery is a cognitive function that involves a distributed cortical network and high-level feedback, but no corresponding feedforward input from the eye, mental imagery-related information should be predominant at cortical depths that contain such feedback. In particular, as participants were instructed to imagine a stimulus centrally, we expected decoding to be above-chance level in foveal V1. This was indeed the case: SVM classification could decode the imagined colour only in the central ROI; this information was available at the deepest depth (*p_adj_*=.02, bootstrapped and FDR-corrected; **Fig. 2A**) but not at any other depths (all *p_adj_*>.05). There was no significant decoding in any of the depth layers in the peripheral V1 region (all *p_adj_*>.05; **Fig. 2B**), which was at the fringes of the location of imagery.

**Figure 2.**
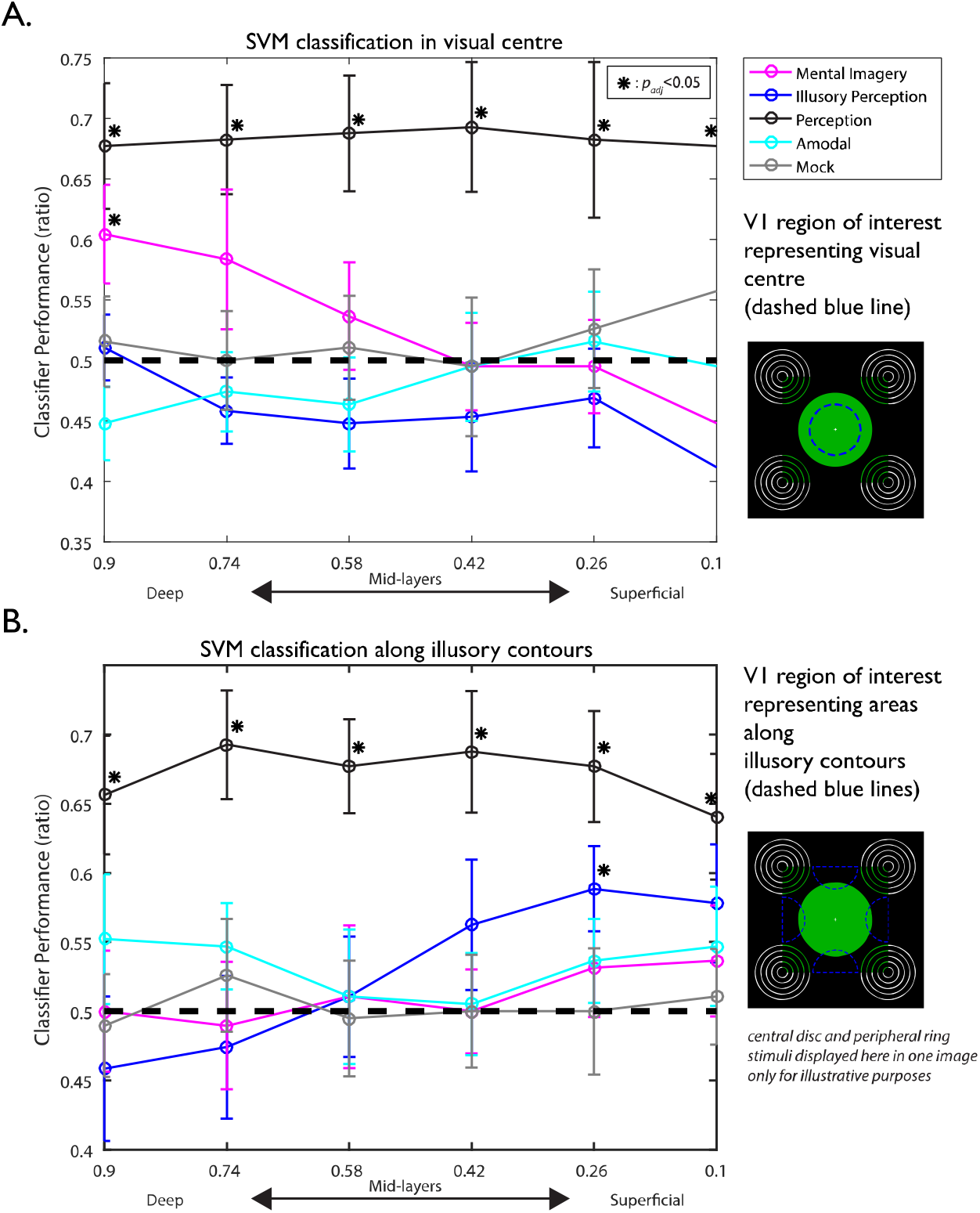
SVM decoding results. SVM decoding results in the V1 region that (**A**) represents the central visual field, and (**B**) the peripheral portions along the illusory boundaries (dashed blue lines in the images on the right). The black dashed lines represent chance level; asterisks denote significant above-chance decoding (*p*_adj_<.05); error bars represent ±SEM. Also see **Supplementary Fig. 8** and **9** for individual decoding plots of each subject. When participants imagined a central disc, colour was decoded only in the deepest depth layer (pink line). Importantly, this finding was restricted to the foveal ROI; no significant decoding was found in the peripheral ROI, which lay at the fringes of the imagery field. Conversely, we could only decode the illusory colour (blue line) in the peripheral ROI, which represented areas along the illusory boundaries of the neon colour spreading stimulus. Here, significant decoding of illusory colour was restricted to superficial depths. In the perceptual condition (black line), we could significantly decode colour at all depths, in both the foveal and the peripheral ROI. In the two control conditions of the illusory stimulus, we found no significant decoding in either of the two ROIs (grey and cyan lines).

Perceptual illusions like the neon colour spreading illusion are thought to arise because higher-order visual areas with larger receptive fields assume the presence of a shape. Feedback from nearby visual areas travel through and arrive in superficial layers(*13*). Previous fMRI research has indeed shown that information about the contextual surround can only be decoded in the superficial layers(*18*). However, another study found an increase in fMRI activity levels during illusory perception that was limited to the deeper layers(*9*). We only found significant decoding at the second most superficial depth (*p_adj_*=.007; also see **Supplementary Fig. 4** & Methods), and this relationship was confined to the V1 regions representing the areas along the illusory contours, where the illusion is most vivid.

Perception is governed by both feedforward and feedback processes, and should be decodable across all layers(*18*). This was indeed what we found: physical colour was decodable across all layers in the V1 region that represented the central visual field (all *p_adj_*<.02). The same was found for the V1 region that represented the peripheral areas at the fringes of the coloured disc (all *p_adj_*<.01).

In the ‘amodal’ version of the neon colour spreading illusion, where the illusion is attenuated by a superimposed white contour, we did not find any significant above-chance decoding in either of the ROIs (all *p_adj_*>.23), nor for the mock version of this illusion, where no illusion should arise (all *p_adj_*>.3).

The pattern of significant decoding seems to reflect the conscious visual experience in space: in the centre, where our participants imagined the colour, we could decode imagery. In contrast, the illusion was weak here, and illusory colour was not decodable. Conversely, in the visual periphery, where the illusory boundaries were located, but which comprised only a small portion of the imagery area at its fringes, we found the opposite pattern. Here, we could decode the illusion, but we could not decode imagery. In addition, the physical stimulus provided during the perceptual condition was decodable across both regions, both in the centre and in the peripheral V1 region, which captured only the fringes of the central coloured disc. To corroborate these observations, we computed a linear mixed effects model with a 2 (foveal vs. peripheral V1 region) x 5 (stimulus conditions) x 6 (cortical depth) design. Among the 3 main effects, only the main effect of stimulus condition was significant (*χ*^2^(4)=17.33, *p*=.002). Furthermore, in line with our observation that imagery is only decodable in the foveal V1 region, whereas the illusion is only decodable in the peripheral V1 region, we found a highly significant 2-way interaction between V1 region and stimulus condition (*χ*^2^(4)=21.34, p<.001). The 3-way interaction between V1 region, stimulus condition and depth did not reach significance (*χ*^2^(20)=29.23, *p*=.08).

Due to larger blood vessels at the pial surface of the cortex, the signal of gradient-echo fMRI imaging is stronger in superficial depths(*19*). This effect was also present in our data (**Supplementary Fig. 5-7**). On the one hand, one could argue that SVM classification performance could be influenced by this, in the way that decoding accuracy could be higher when the overall signal – and hence the signal-to-noise ratio – is enhanced. On the other hand, the fMRI signal might also be more spatially specific when the point spread of the blood-ogygen-level-dependent (BOLD) signal is smaller due to smaller vessels at deeper depths(*20*). This, in turn, could lead to better decoding at deeper depths. The pattern of our results however does not suggest that SVM classification is dictated by overall BOLD activity levels at different cortical depths: we found that imagery was only decodable in the deep layers, where the signal strength was the lowest; conversely, illusory perception was only decodable in the superficial layers, where the signal was highest; and the perception of physical stimuli was decodable at all depths, regardless of different signal strengths.

Taken together, we find that the laminar representation of low-level feedback in V1 during illusory perception is distinct to that of high-level processes like mental imagery. We still need to better understand the precise connectivity architecture to V1 layers, as well as the origins of the projections. Our findings suggest that feedback to the deep and superficial layers may serve different functions: Low-level feedback to superficial layers may be linked to perceptual or perception-like experiences that appear embedded in our sensory environment. In contrast, feedback to deep layers may serve the formation of a more malleable ‘inner’ sensory world, which we can keep separate to our ongoing perception. This difference could also be a decisive factor that sets apart normal everyday imagery from pathological forms of uncontrollable, seemingly ‘real’ imagery, e.g. in Parkinson disease(*21*) or the Charles Bonnet syndrome(*22*). More research will also be needed to disentangle whether our ability to exert volitional control over conscious sensory experiences and our perception of ‘realness’ are linked, or independent of each other. Our finding that mental imagery is only decodable in specific V1 layers will further help us to refine theories that mental imagery is ‘perception-like’ in nature(*12*): Unlike perception, the processing of imagery appears to be confined to certain layers of V1. This important difference had been missed with conventional fMRI technology, which lacks the precision to look at brain signals at different cortical depths.

## Acknowledgements

This project has received funding from the European Union’s Horizon 2020 Framework Programme for Research and Innovation under the Specific Grant Agreement No. 720270 and 785907 (Human Brain Project SGA1 and SGA2). We thank Lucy Petro for suggesting to add a condition with an amodal version of the neon colour spreading stimulus. We also thank Jane Alfred for helpful comments on the manuscript.

## Author contributions

J.B. and L.M. independently conceived the study, and then designed the study together. J.B. acquired, pre-processed, and analysed the data. A.M. computed the visual projections. J.B. wrote the manuscript, and all authors edited the manuscript.

## Supplementary Materials

### Methods

#### Participants

fMRI data were collected from 18 healthy participants with normal or corrected-to-normal vision (age range 19-36 years at the time of the experiment, M=23.3, SD=4.96; 13 females and 5 males). These participants were recruited from a larger participant sample of 55 individuals (age range 18-36 years; 20 males, 35 females), who underwent a behavioural pretest designed to quantify individual imagery strength (see next section). Of this pre-sample, 21 participants had an imagery strength score that met or surpassed an a priori defined threshold, and were invited to take part in the main fMRI experiment, 18 of whom completed the fMRI experiment. Due to bad inter-run alignment in two data sets (see functional GE-EPI data preprocessing), analyses were conducted with the data sets of 16 participants. Participants were recruited from the participant pool of the Institute of Neuroscience and Psychology, University of Glasgow. Preceding the study, fMRI piloting sessions were conducted with 3 participants at the Maastricht Brain Imaging Centre, Netherlands. All subjects gave informed consent and the study was approved by the ethics committee of the College of Science and Engineering of the University of Glasgow.

#### Behavioural pretest to quantify individual imagery strength

The ability to imagine visual content varies largely within the population(*23–26*). Furthermore, an individual’s ability to form a mental image is associated with enhanced decoding accuracy of fMRI activity patterns in early visual cortex(*27*). This suggests a link between mental imagery ability and the precision of imagery-related visual cortex signals measured by fMRI. To increase the probability of finding meaningful imagery–related V1 activity patterns and thereby increase the statistical power of the study(*28*), we ran behavioural pre-screenings to identify individuals with good visual imagery abilities. To do so, we used a behavioural paradigm that quantifies individual imagery by measuring its impact on subsequent conscious perception of a binocular rivalry display(*11, 24–26*). Binocular rivalry occurs when the two eyes see two different images, one to each eye. This setup results in the phenomenon that perception alternates between the two images, with one image reaching conscious awareness while the other one is suppressed(*29, 30*). Importantly, previous work has shown that preceding imagery can bias brief subsequent binocular rivalry perception, with the previously imagined stimulus having a higher chance to gain dominance. This bias is further increased with longer periods of imagery(*11*), and when the stimulus is imagined more vividly in a given trial(*26, 31*). In addition, individual imagery ability quantified by this paradigm is linked to individual visual working memory storage(*24, 25*), and is highly stable over time(*26*). Taken together, the evidence suggests that determining individual imagery ability by quantifying its impact on subsequent binocular rivalry perception might be a valid and reliable method with which to identify individuals with good imagery ability.

In the behavioural pre-test session, participants sat in a darkened room at a distance of 60 cm from a computer screen (ASOS PG278Q, 27”), their heads stabilized using a chin rest. They wore crystallized shutter glasses (NVIDIA 3D VISION 2), which allows different images to be projected separately to each eye, thereby making it possible to induce binocular rivalry. Experimental stimulation was controlled using MATLAB R2016a, in combination with the Psychtoolbox v3.0.13 extension(*32*), running on a DELL Precision T3500 computer with an Intel^®^ Xeon^®^ CPU processor and an NVIDIA GeForce GTX 970 graphics card. Participants were instructed to maintain fixation on the central fixation cross throughout the experiment. Apart from the white fixation cross, the screen remained black. Participants completed two tasks. Prior to the imagery strength task, participants did an eye dominance task designed to adjust the luminance of the two stimuli to each individual’s eye dominance. Individual differences in eye dominance would otherwise affect binocular rivalry perception, resulting in a bias for the image that is viewed with the more dominant eye. The procedure has been described in detail previously(*11, 26*); in short, it applies an adaptive testing procedure designed to individually adjust the luminance of the two colours such that it is equally likely for each to gain perceptual dominance. Following the eye dominance task, participants completed a couple of training trials of the imagery test to familiarize themselves with the task. If necessary, further adjustments of the luminance values were done before starting the main imagery task.

At the beginning of each trial of the imagery strength task (**Supplementary Fig. 1A**), a grey letter ‘R’ or ‘G’ (1.3cm in size, i.e. 1.2° visual angle) appeared in the central lower part of the screen, at a distance of 4.9 cm (4.7°) from the fixation cross. The letter cued the participants as to which of the two colours they should imagine in the given trial – ‘R’ for red, ‘G’ for green. The cue was shown for 1 sec, and was followed by a 1 sec break, during which only the fixation cross was visible. Following this phase, a faint grey circle (radius of 3.65cm; 3.5°) was presented for 7 sec, centred on the fixation cross. During these 7 seconds, participants were asked to imagine the colour as vividly as possible and within the frames of the faint grey circle on the screen. Following this 7-second imagery phase, the circle disappeared and the word ‘vividness?’ appeared in the central lower part of the screen (at a distance of 4.9 cm/4.7° from the fixation cross), instructing participants to rate, on a scale from 1 to 4, how vivid their mental image of the colour had been. 1 represented the lowest and 4 the highest level of vividness. After responding with a key press, the binocular rivalry display appeared for 0.75 seconds, showing circular Gaussian-windowed green and red colour stimuli, one shown to each eye. When the binocular rivalry display disappeared, participants were instructed to indicate via a key press which of the two colours had been dominant (‘1’ for red, ‘2’ green, ‘3’ for mixed). 10% of the trials were catch trials, where a mixture of both colours was presented to each eye; in these trials, participants should give a ‘mixed’ response; the failure to do so is an indicator of decision bias. Each participant completed one run with n=100 trials, with one break after completing half of the trials. Individual imagery strength was defined as the individual bias with which the imagined stimulus gained perceptual dominance during subsequent binocular rivalry: imagery strength (%primed) = n_primed_/(n-n_mock_-n_mixed_), with n_primed_ being the number of (non-catch) trials in which the imagined colour matched the colour that was subsequently dominant during binocular rivalry, n the total number of trials, n_mock_ the number of catch trials, and n_mixed_ the number of (non-catch) trials in which participants reported a mixed percept. An arbitrary, *a priori*-defined threshold of 60% priming was set as a threshold to identify individuals with good enough imagery ability (**Supplementary Fig 1B**). Individuals that scored priming values at or above the threshold level were invited to participate in the fMRI part of the study. The subjective vividness ratings that participants gave were not used as a criterion for participation in the fMRI sessions, but were used in further analyses to check the validity of the pre-test (**Supplementary Fig 1C and 1D**).

#### fMRI experimental procedure and design

In a 7T-MRI scanner, participants completed 6 runs of task-related experimental stimulation, and 2 retinotopic mapping runs. Throughout the experiment, participants were asked to maintain fixation on the white central fixation cross on an otherwise black screen. Fixation was monitored using an eye tracker (EyeLink 1000 Plus, SR Research). However, the eye tracker was often not able to track the eyes accurately throughout the runs because the head coil limited the camera’s field of view. This rendered the data unusable for most participants. The screen was attached to the top end of the scanning table and viewed through a mirror attached to the head coil, at a viewing distance of approx. 96 cm. The task consisted of 5 conditions, in which participants were asked to view or imagine coloured shapes that were either red or green (**Fig. 2A**). Each condition was shown in blocks of 8 seconds, interleaved by inter-stimulus intervals (ISIs) that were jittered between 7 and 8 seconds. In each run, each condition was shown 6 times, i.e. 3 times with red stimuli, and 3 times with green stimuli. In the perception condition, participants viewed a uniformly red or green disk presented at the centre (radius of 3.1°; in 4 participants, the radius was 3.7°), flashing at a frequency of 4 Hz. In the imagery condition, participants were asked to imagine that same red or green disc, with the area of imagery indicated by a faint grey circle around the fixation cross. The letter ‘R’ or ‘G’ cued participants as to which colour to imagine (red or green). It was presented for 1 sec, 2 secs prior to the presentation of the faint grey circle, which signaled the onset of the imagery task. Importantly, the cue was presented in the periphery outside of the grey circle presented subsequently, and its location on the screen was randomized, but kept at a stable distance of 4.3° to the central fixation cross (5.1° in 4 participants). As some participants had difficulties with seeing the lower central part of the screen due to the head coil, this part of the screen was spared (i.e. the cue never appeared here). In the illusory perception condition, participants viewed a visual illusion, known as neon colour spreading, in which 4 peripheral pacman-shaped stimuli elicit the illusion of a coloured (red or green) square, with illusory coloured boundaries continuing beyond the edges of the coloured pacman shapes in the periphery. The four pacman-shaped stimuli were each presented in one of the four quadrants of the screen, respectively, with the centres of the stimuli located at a distance of 6.24° to the central fixation cross (7° in 4 participants). Each consisted of 5 concentric rings of increasing radii, the distance between radii remaining constant. The outermost (and largest) ring’s closest distance to the central fixation cross was 3.94° (4.71° in 4 participants). The inwardly directed quarter of the rings was either red or green, thereby leading to the illusion of a coloured square. The remaining three quarters of the ring were white. In the third condition, we presented an ‘amodal’ version of this illusion, in which a white contour was placed in the area between the pacman-like rings, thereby attenuating the illusion (the contour’s closest distance to the central fixation cross being 3.86°; 4.61° in 4 participants). In the mock version of the illusory perception condition, the pacmans were rotated outwards, such that no illusion of a coloured shape should arise. At the end of each run, we also presented contrast-inverting (4 Hz) checkerboard patterns. One was a disc in the central target region with a radius of 2.1° (2.6° in 4 participants) from the central fixation cross, and one was ring-shaped, surrounding the central target region (spanning an area from a radius of 2.1° to 4.4° from central fixation; 2.6° to 4.9° in 4 participants). The target > surround contrast was used to map the central target region in V1, and to compare this region with the central target region identified by population-receptive field (pRF) mapping to visually crosscheck the validity of the pRF mapping (see below). The ROI definition was then made on the basis of pRF mapping. Note that this central target region identified by the target vs. surround contrast and by the pRF mapping was smaller than the area in which the stimulus was viewed or imagined in the perception and imagery condition. We did this for two reasons: First, to ensure that even with small shifts of the centre of gaze due to unavoidable eye saccades, the coloured disc would still be represented by foveal V1 neurons. Second, it limited the risk that despite small gaze shifts, the peripheral pacman stimuli of the illusory and control conditions would not fall within the receptive fields of the foveal V1 region that we decoded from, as the pacmans contained actual (and not just illusory) colour. Following the 6 task-related runs, we also acquired Polar Angle and Eccentricity maps. These were used: (1) to map V1 retinotopically on the cortical surface(*33, 34*); and (2) to compute pRF maps, to determine which portions of V1 represent which portions of the visual field(*17*). In the polar angle mapping run, participants viewed a contrast-inverting (5 Hz) black and white checkerboard wedge rotating anti-clockwise around the central fixation cross. The wedge had an angle of 22.5° and rotated around the fixation cross 12 times, each cycle lasting 64 seconds. In the Eccentricity mapping run, participants viewed a contrast-inverting black and white checkerboard ring, slowly expanding from the centre of gaze to the periphery. An expansion cycle lasted 64 seconds, and was repeated 8 times. The Polar Angle and Eccentricity measurement induces phase-encoded neural activity that allows us to estimate the boundaries between early visual areas (Polar Angle) and to match the different expansion radii of the visual stimulus to the eccentricity radii on the cortical surface (Eccentricity mapping). Both wedge and ring stimuli were presented in front of a grey background with a target spanning the screen, centred on the fixation cross. Before and after the presentation of the wedge or the ring, there was a baseline phase of 12 seconds during which only the background and fixation cross were presented. In both polar angle and eccentricity runs, the participants’ task was to maintain fixation on the central fixation cross.

#### fMRI data acquisition

fMRI data were acquired in a Siemens 7-Tesla Terra Magnetom MRI scanner with a 32-channel head coil located at the Imaging Centre of Excellence of the University of Glasgow. Each of the 6 task-related runs consisted of 272 T2*-weighted gradient-echo echoplanar (EPI) images using the CMRR MB sequence with an MB factor of 1 (voxel resolution: 0.8mm^3^ isotropic resolution, distance factor: 0%, 27 slices, FoV=148mm, TR= 2000 ms, TE=26.4 ms, flip angle: 70°, slice timing: interleaved, bandwidth = 1034Hz/px, phase encoding direction: head to foot). We used the same sequence for the retinotopic mapping runs, acquiring 396 volumes for the Polar Angle and 268 volumes for the Eccentricity mappings. In order to identify susceptibility-induced distortions, which can distort the functional EPI images and make it hard to align them to the anatomical images, we also recorded 5 volumes of the same EPI sequence with the phase-encoding direction inverted. This way, it is possible to estimate field distortions and correct for them using FSL’s topup tool if necessary(*35, 36*). In addition, whole-brain, high-resolution T1-weighted MP2RAGE images were acquired (voxel resolution: 0.63mm^3^ isotropic resolution, 256 sagittal slices, TR = 4.68 ms, TE = 0.00209, TI_1_ = 0.84ms, TI_2_ = TI_2_=2.37, flip angle_1_ = 5, flip angle_2_ = 6, bandwidth = 250Hz/px, acceleration factor = 3 in primary phase encoding direction, FOV = 240mm). In order to scale the applied voltage to achieve accurate flip angles for the scanning session, we also ran a 3DREAM B1 mapping sequence beforehand(*37*) (voxel resolution: 4.5mm^3^ isotropic resolution, 44 slices, FOV = 288mm, TR = 5000ms, TE_1_= 0.9ms, TE_2_ = 1.54ms, flip angle_1_: 60°, flip angle_2_: 8°).

#### functional GE-EPI data preprocessing

Functional imaging data were analysed using BrainVoyager 20.6(*38, 39*). Functional image preprocessing involved 3D-motion correction, slice time correction, high-pass filtering, and coregistration to the T1-weighted anatomical images. To ensure the functional runs were excellently aligned to each other, which is imperative for meaningful multivariate pattern analyses with high-resolution voxels, we computed inter-run spatial cross-correlations of the signal intensities of the functional volumes. Where necessary, Brainvoyager’s VTC-VTC grid search alignment and spatial transformation tools were used to improve inter-run alignment until it was at least r>.9 on average. Further, functional-anatomical alignments were checked visually to ensure that the functional scans were well aligned to the anatomical image at the location of the ROI. Due to the relatively small ROIs, no susceptibility-induced distortion correction using FSL’s topup tool was necessary. However, the data of 2 of the 18 participants showed very bad inter-run alignment (r=.13 and r=.38, respectively), and after several failed attempts to improve the alignment, the two data sets were removed from further analysis.

#### T1-weighted anatomical data preprocessing

We first processed the anatomical imaging data with a range of brain imaging software, before converting them to BrainVoyager format to continue processing. We used FSL to upsample the anatomical data to 0.4mm(*40*) (FMRIB’s Software Library, www.fmrib.ox.ac.uk/fsl). For a preliminary white matter-gray matter segmentation, we then processed anatomical imaging data using a custom-written pipeline described previously(*41, 42*). The pipeline uses R(*43*), AFNI(*44*), and nighres(*45*). Where necessary, we corrected the results of the automatic segmentation manually using ITK-SNAP(*46*). Our focus of the manual corrections was on V1, and we took great care to ensure that the grey matter segmentation result did not contain any parts from the sinus or skull. The corrected segmentation was then used to mask the original images using FSL. We then converted the masked anatomical image, as well as the white matter mask, to Brainvoyager format to continue processing in Brainvoyager. After a rigid-body transformation of the anatomical images into ACPC space, the white matter mask was drawn onto the anatomical image in a first step, before using Brainvoyager’s advanced segmentation tool for the GM-CSF segmentation. From the segmented white matter-grey matter boundary, we reconstructed the surface of the occipital lobe. The retinotopic mapping-based estimation of V1 boundaries and the definition of the ROIs within V1 was then done on the inflated cortical surface (see below). This process was done individually for every data set.

#### Retinotopic mapping of V1 boundaries

We estimated the boundaries of V1 using the fMRI data recorded during the previously described retinotopic mapping scans. In order to estimate V1 functionally, a Fourier transform was applied to each voxel’s fMRI time series of the polar angle and eccentricity mapping run in order to compute amplitude and phase at stimulation frequency(*33, 34*). The different phase angles were then colour-encoded and mapped onto the inflated cortical surface. Each colour represented an F-ratio of the squared amplitude divided by the average squared amplitudes at all other frequencies. On the basis of the colour encoding, the boundary segregating V1 from V2 was then estimated manually on the cortical surface for each individual subject. Due to the small slab of the functional volume, only V1 could be estimated. In 2 of the 16 participants, the colour encoding did not reveal a retinotopic map in one of the hemispheres (left hemisphere in S12 and right hemisphere in S13), and therefore V1 could only be estimated in the other hemisphere.

#### Regions of interest definition in V1 using population receptive field mapping (pRFs)

Following the retinopic mapping-based estimation of V1, we identified our regions of interest in V1 using an approach informed by pRF mapping (**Supplementary Fig. 2**). Using visual field mapping stimulation, population receptive field mapping allows to estimate which portion of the visual field an fMRI voxel is most responsive to(*17*). To compute pRFs, we first created model time courses that predicted how a voxel’s timecourse responsive to a certain portion of the visual field would look during eccentricity and polar angle mapping stimulation. We assumed a standard Gaussian model; we first created a Gaussian window for every portion of the visual field, defined by its spatial coordinates x and y, as well as by its size (i.e. standard deviation in the Gaussian model). Models of 24 different sizes (i.e. standard deviations) were estimated, with the centre of each Gaussian window being one standard deviation apart from the next. Then, we estimated for each of these models how a voxel’s response to the polar angle and eccentricity stimulation should look over time, if it was responsive to that visual portion. In a next step, we computed the correlations between every V1 voxel’s actual time course, and the predicted time courses for all portions of the visual field. For every voxel, we then determined which visual portion it was most responsive to, based on which model explained most of the variance of the voxel’s responses (*r*^2^) over time.

Using this approach, we identified preliminary ROIs by determining those fMRI voxels whose population receptive fields (with a spread of 1 standard deviation) fell within those portions of the visual field that we were interested in (**Fig. 2B**). To shield from excessive levels of noise in our models, we thresholded *r*^2^, such that only those voxels whose best model’s fit exceeded the threshold were included. This procedure is very conservative and protects against noise, but it also, as a consequence, mostly identifies voxels in the superficial portions of grey matter, which thus do not represent all layers (see **Supplementary Fig. 3** for an illustration of the effect of different *r*^2^ thresholds). This is likely because GE-EPI data generally yield stronger responses in the superficial depths compared to voxels in deeper portions(*19*). To ensure that all cortical depth layers were represented in the ROI, and to correct for the fragmented nature of voxel identification, we used the following procedure. First, we projected the functional voxels in question onto the inflated cortical surface. Second, we manually drew boundaries around the patches and labelled them. Last, we projected these patches back into volume space in order to proceed with the cortical depth layer segmentation. It is known that neurons are organized in hypercolumns(*47*), which means that neurons in the deeper layers represent the same visual portion as those ‘stacked’ on top of them in the more-superficial layers. As a consequence, it is neuroanatomically justified to include voxels from all layers for a portion on the cortical surface that has been identified as a ROI. Overlaid retinotopic maps, and the ROI’s anatomical position on the cortical surface, were also used for additional validation of our approach. Furthermore, we conducted cross-checks using lower thresholds for *r*^2^ to confirm that our approach was adequate. As shown in **Supplementary Fig. 3**, the pRF-defined regions grew into the deeper grey matter portions in question when the *r*^2^-threshold was lowered.

In each individual, we identified 2 ROIs: first, a foveal ROI to examine the information content of the V1 portion that represents the central visual field. This ROI was computed from the same visual portion that was mapped using the target vs. surround contrast from the task-related experimental runs. It spanned a radius of 2.1° from central fixation (2.6° in 4 participants). As a “sanity check”, we visually cross-checked the location of the central ROI mapped by pRFs with the one estimated by the target vs. surround contrast. Second, we determined a peripheral ROI, located at the four illusory boundaries of the illusory square. The visual portions presented four semi-circles, with the straight edge positioned along the illusory boundary (at a distance of 4.2° to the central fixation cross; 4.7° in four participants). Due to the headcoil in the fMRI scanner, which limited visibility of the lower (and, sometimes, upper) central part of the screen in most participants, the majority of the identified voxels had their pRF in the right or left semi-circles, and fewer were located in the lower and upper ones.

#### Cortical depth-layer segmentation

After creating a cortical thickness map, BrainVoyager’s high-resolution cortex depth grid sampling tool was used to segment the grey matter of the ROIs into 6 equidistant cortical depth layers (at 0.1, 0.26, 0.42, 0.58, 0.74, and 0.9 depth). Note that with the coarse resolution of high-resolution MRI compared to histological methods, cortical depth sampling based on equidistance sampling results in only slight differences compared to equivolume sampling(*48*). The comparatively coarse resolution also has the consequence that grids at specified depth levels partly use the same voxels as the neighbouring grids, which means that they are not fully independent. For example, some voxels that were included in the grid at 0.9 depth may also have been part of the grid at 0.74 depth.

#### Univariate and multivariate analyses of ROI activity

To investigate univariate differences across the 6 cortical depth layers of the ROIs in the different conditions, we computed a GLM analysis, modelling each condition with two-gamma hemodynamic response functions (HRF), time-locked to the presentation of the stimuli in the experimental runs. The two colours in each condition were modelled separately, resulting in a total of 10 factors of interest. The target and surround mapping block at the end of the experimental runs were regressed out as additional factors.

For the multivariate pattern analysis, a linear SVM classification with default parameters was computed. Voxels whose raw mean BOLD signal intensities were below 100 were removed prior to the analysis. Beta weights for every trial were then estimated in a GLM analysis. Before entering the data into the SVM classifier, the beta weights were normalized by rescaling the values between −1 and 1. Cross-validation was performed in a leave-one-run-out manner. The reported SVM accuracies was the average across the cross-validation folds.

Both the univariate and multivariate analyses were conducted using custom-written scripts in MATLAB R2015b & R2016 in connection with BVQX v0.8b /NeuroElf v0.9c toolboxes (http://neuroelf.net/). The linear SVM classification was computed using the LIBSVM toolbox(*49*), v2.86.

#### Illusory perception vs. mock illusion decoding and visual projections

Previous work has found that illusory contours involve a selective activation of deep V1 layers(*9*). In contrast, in our multivariate pattern analysis, we can only decode illusory colour at superficial depths in our study. However, differences in the analyses of these two studies could have contributed to these differences in findings. In our study, we decoded two illusory colours against each other, whereas Kok et al. looked at (univariate) activity differences between the condition when an illusory figure was present and when it is was not. For better comparability with Kok et al.’s results, we ran an additional SVM classification analysis, where we decoded the illusory perception condition against the mock version of the illusion. As the stimuli in our study were presented in two different colours, we could compute SVM classification accuracies across the 6 cortical depths twice, once for the red stimuli, and once for the green stimuli. This also allowed us to assess the reliability of the analysis. As shown in **Supplementary Fig. 6A**, decoding accuracies increased towards the more superficial layers in both colours; the red stimuli showed significant decoding in more layers than the green stimuli, but both red and green stimuli showed significant above-threshold decoding in the second and third most superficial depth (both *p_adj_*≤0.001). This is not in line Kok’s et al.’s results, who found selective deep layer involvement, but is in line with our finding that illusory colour can only be decoded at the second superficial depth, as reported in the main text. It is also supported by findings in nonhuman primates(*50*) and mice(*51*) that edges between figure and ground lead to additional V1 activity caused by synaptic input into layer 2/3. In addition to computing SVM classification accuracies, we also projected voxel influence on SVM classification(*52*) from the different cortical layers into visual space using a weighted average method based on voxel pRF models(*53*) (see **Supplementary Fig. 6B**). Instead of displaying voxel influence in cortical space, this approach thus translates voxel influence into visual space, giving an idea of which visual field portions (or pixels) contribute most to SVM classification. Here, we were interested in which sub-portions of the visual field areas represented in the peripheral V1 ROI were particularly influential for SVM classification. The method takes the following form:

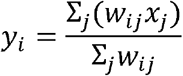

where *y_i_* is the activity at a visual field pixel *i*, and *x* is a vector of SVM weights converted to voxel activity. Note that the term *activity* here does not refer to BOLD activity, but to the influence that different portions of the visual field have on SVM classification analyses. *w* is a matrix of vectorised pRF functions for the voxels in *x*, with columns *j* representing voxels, and rows *i* representing how much a particular visual field pixel is represented by a particular voxel. *y_i_* can become arbitrarily inflated when Σ_*j*_*w_ij_* is extremely small, as is the case in areas of the visual field without sufficient pRF coverage. We therefore truncated each pRF function to have only weights within 2σ of its centre and defined *y_i_* = 0 for pixels with Σ_*j*_*w_ij_* = 0.

#### Statistical testing

To test the mean SVM classification accuracy across subjects for each stimulus condition at each cortical depth against chance level (50%), we used bootstrapping with 10,000 samples. The statistical significance level of α=0.05 was corrected for multiple comparisons using false discovery rate (FDR; 60 comparisons across all 6 layers, 5 conditions and 2 regions of interest). Further, to test the main effects and interaction effects between V1 region, stimulation condition and cortical depth, we computed a 2 (foveal vs peripheral ROI) x 5 (stimulus condition) x 6 (cortical depth) Linear Mixed Effects Model using R(*43*) with lme4(*54*). V1 region, stimulus condition and cortical depth were modelled as fixed effects. As random effects, we modelled intercepts for subjects, as well as by-subject random slopes for the effects of V1 region, stimulus condition and cortical depth. *χ*^2^ and *p*-values of the fixed effect were estimated using maximum likelihood ratio tests of the full model, which contained the effect in question, against the reduced model which did not contain the effect. We inspected the plots of the residuals visually and computed Shapiro-Wilk tests to ensure they did not show deviations from homoescadicity and normality.

To analyse the relationship between imagery strength and subjective vividness in the behavioural pre-test data, we used Spearman rank correlation for the between-subject analysis (**Supplementary Fig. 1C**), as the normality assumption for the imagery strength values was violated (Shapiro-Wilk normality test: W=0.90, p<.001). Further, to examine the trial-by-trial relationship between imagery-induced priming and vividness, we computed a linear mixed effects model with a 2 (group) x 4 (vividness rating) design (**Supplementary Fig. 1D**). As a random effect, intercepts for subjects were modelled. As the residuals were not normally distributed (*W*=0.96, *p*<.001), an ordered quantile normalization was applied to transform the data before computing the model.

In addition, we tested the main and interaction effects of the univariate analysis of the fMRI activity in the two ROIs with a 2 (foveal vs peripheral ROI) x 10 (stimulus conditions, separate for each colour) x 6 (cortical depth) design. As the residuals were still not normally distributed (*W*=0.96, *p*<.001), after using bestNormalize(*55*) to find the best transform of the data, a generalized linear mixed effects model was used instead of a linear mixed effects model. As random effects, we modelled intercepts for subjects, as well as by-subject random slopes for the effects of V1 region, stimulus condition and cortical depth. A Gaussian distribution was used as a GLM family, with the log-function as the model link function.

## Supplementary figures

**Supplementary Figure 1.**
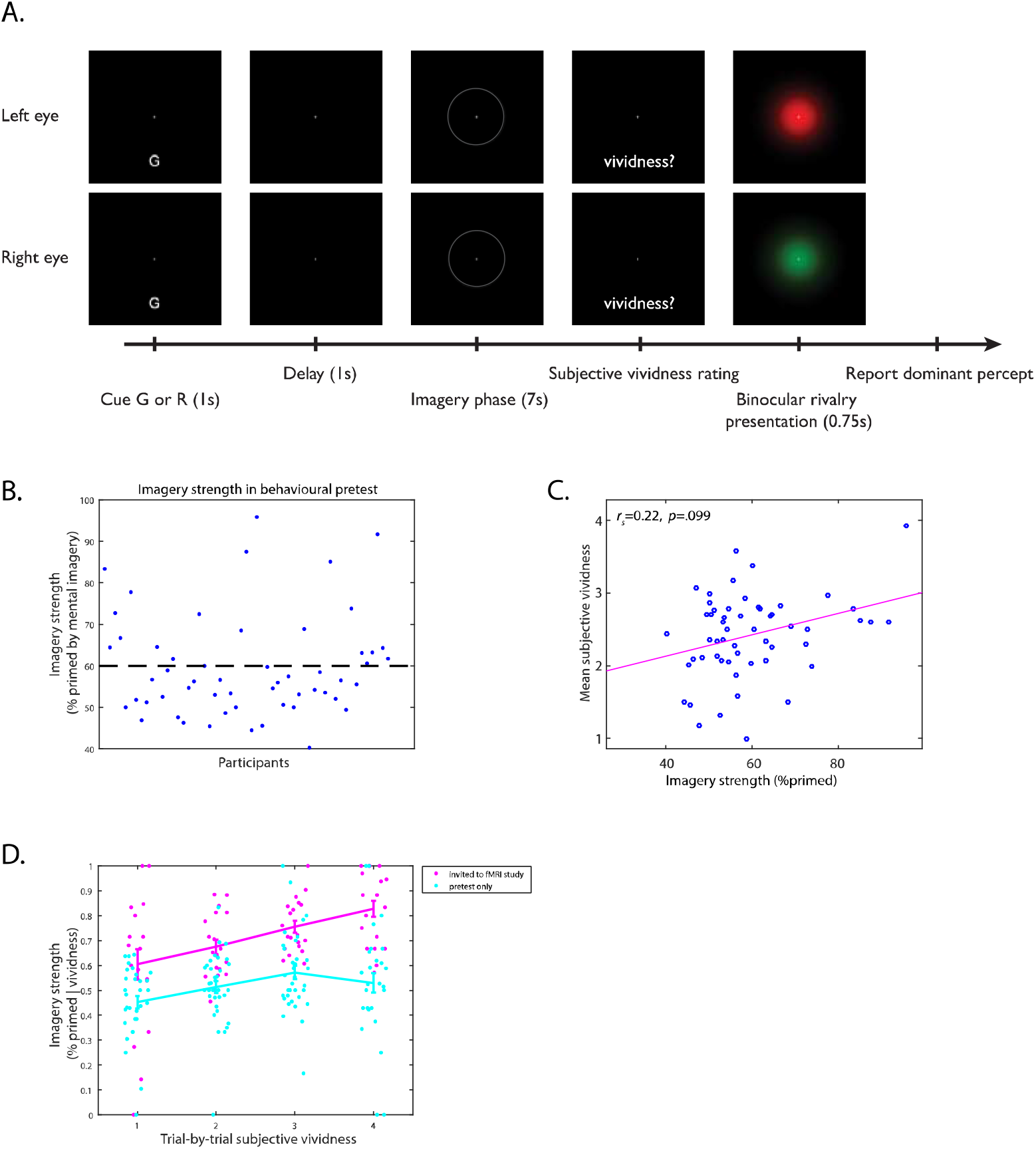
Behavioural pretest procedure and analysis. **A.** Experimental procedure. At the start of each trial, a letter cue indicated which colour to imagine. Following a short delay, participants had 7 seconds to imagine the colour. A faint circle presented in the screen centre around the fixation cross indicated the visual field location within which the participants were asked to imagine the colour. Following the imagery phase, the participants were asked to rate the subjective vividness of the imagined stimulus (on a scale from 1-4, with 1 denoting low vividness, and 4 high vividness). Following this, a binocular rivalry stimulus was flashed, containing Gaussian-windowed images of the two colours, one shown to each eye. When the stimulus disappeared, participants indicated via button press which of the two stimuli had been dominant. **B.** Only participants whose imagery strength (imagery-biased %primed) was at or exceeded the *a priori* defined threshold of 60% (dashed line) were invited to participate in the fMRI experiment. Each data point denotes one participant, and the dashed line indicates the 60% threshold. **C.** In line with previous findings (Bergmann et al., 2016), individual mean vividness and imagery strength did not correlate significantly with each other. Each data point denotes the value of one participant; Spearman rank correlation was used. **D.** In contrast to the non-significant correlation between mean vividness and imagery strength on a between-subject level, a relationship between vividness and imagery-induced priming can be observed on a trial-by-trial level. Here, we split up the sample into those participants who were invited for the fMRI part of the study (pink dots), and those whose %primed value was below the cut-off and who were consequently not invited to undergo fMRI (cyan dots). Error bars represent ±SEM. In both groups, a higher vividness rating in a given trial was associated with a higher probability of subsequent priming during binocular rivalry (significant main effect of vividness in a linear mixed effects model, *χ*^2^(3)=22.06, p<0.001). Note that not every participant used the whole scale for their vividness ratings, which results in different amounts of data points per vividness rating. These findings again replicate previous findings (Bergmann et al., 2016).

**Supplementary Figure 2.**
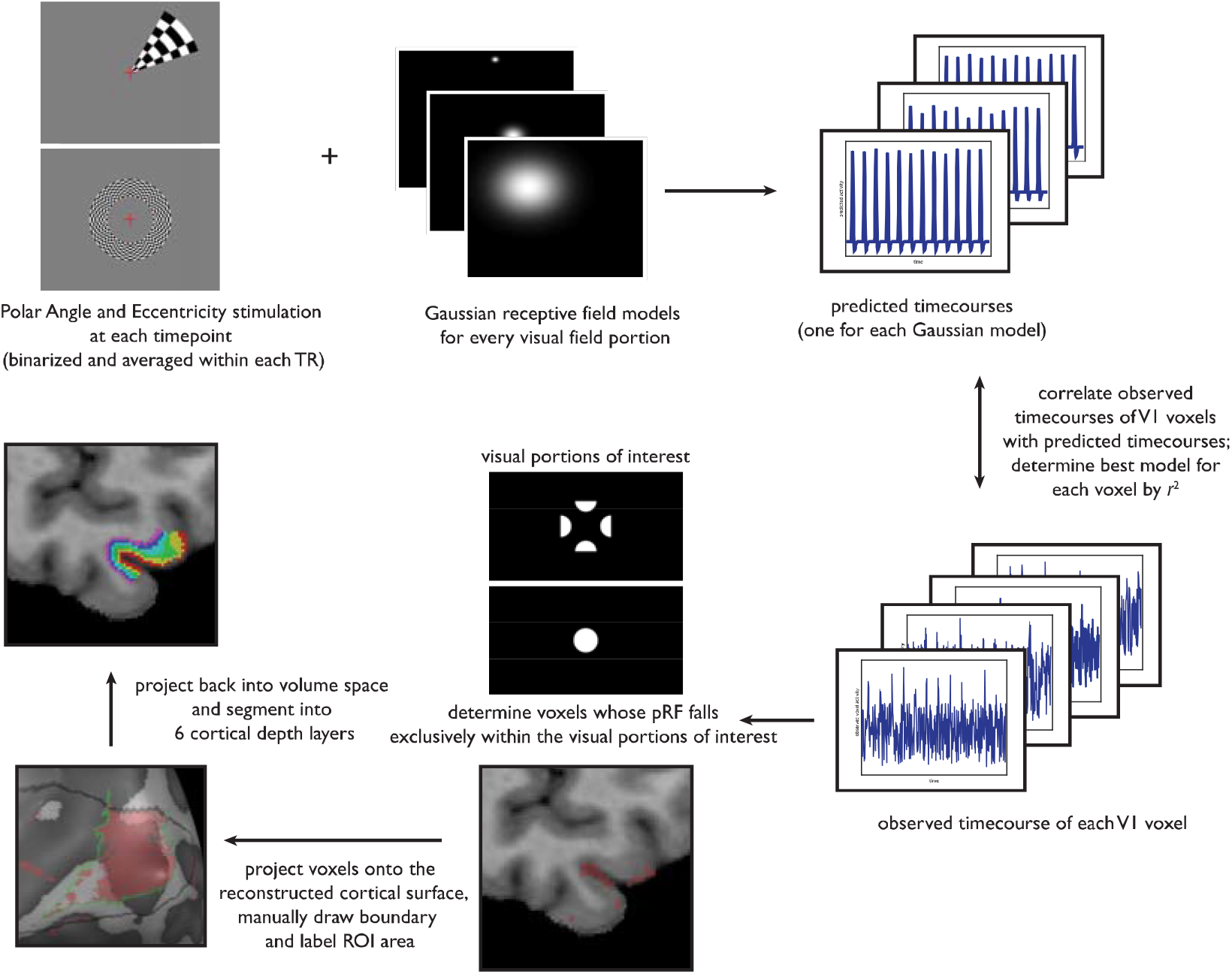
Region-of-interest definition procedure. All participants underwent a retinotopic mapping procedure after the experimental task. This was used to 1) map the boundaries of V1, and 2) estimate population-receptive field (pRF) models. To obtain pRF models, information about the stimulus sequence during Polar Angle and Eccentricity stimulation is needed. For this purpose, we first generated binarized stimulus frames of the Polar Angle and Eccentricity stimulation for each timepoint, which were then averaged within each repetition time (TR). Next, we created Gaussian models of different sizes for every visual field portion. For each Gaussian model, we calculated how a voxel’s response to the retinotopic mapping stimulation should look like over time, if it was responsive to the visual portion modelled by the Gaussian. These models were then correlated with the actual timecourses of V1 voxels. The Gaussian model that explained most of the voxel activity’s variance (*r*^2^) was then selected as best model. With this approach, we could determine which voxels represented those portions of the visual field that we were interested in. To exclude excessive levels of noise, we set a threshold for *r*^2^. The consequence of this is that most of the voxels above the threshold are close to the pial surface, where the overall BOLD activity strength is higher (De Martino et al., 2013). Further cross-checks confirmed the validity of the approach (see **Supplementary Fig. 3** and Methods). The voxels were then projected onto the cortical surface, and after crosschecking the voxels’ location with the anatomy and the retinotopic maps to ensure the models were anatomically accurate, we manually drew boundaries around them and labeled the patches. Following this, the labelled patches of interest were projected back into volume space, where they were segmented into 6 cortical depth layers.

**Supplementary Fig. 3.**
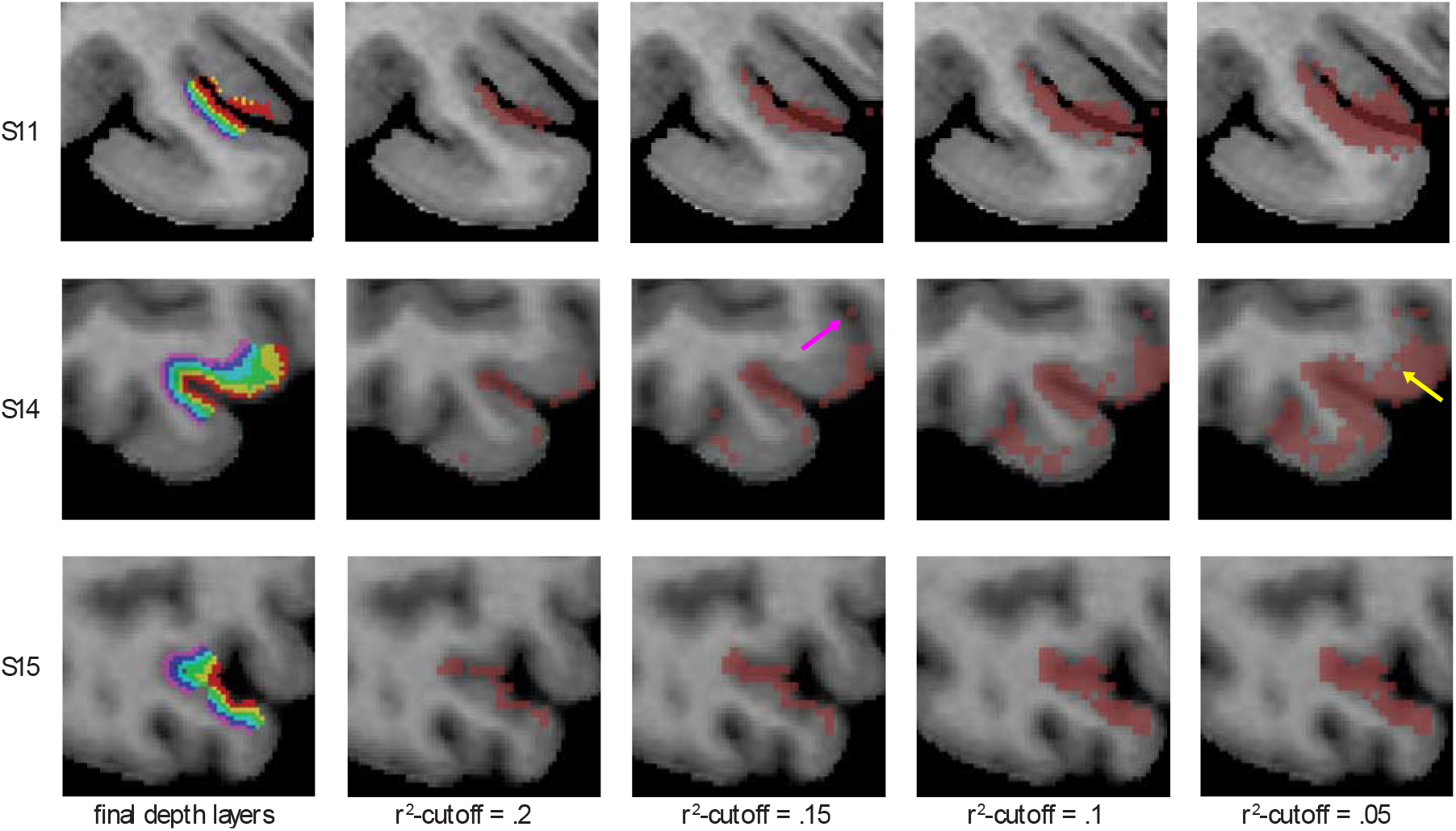
Three example subjects illustrating the influence of different *r*^2^-thresholds on voxel selection. Each row represents one subject. Final cortical depth layers are shown in the left-most column, and preliminary region of interest (ROI) estimates (dark red voxels) with different *r*^2^-thresholds are shown in the four columns to the right (*r*^2^-thresholds are lowered from left to right). Depending on the noise levels of the fMRI signal, population-receptive field models can be heavily influenced by noise. As a result, the best Gaussian model of some voxels is located within the visual areas of interest, even when it is evident from the anatomy that these voxels might represent different portions of the visual field (i.e. they are located far away from the other V1 voxels whose best models also fall within the same visual area of interest; for example, see the voxel indicated by the pink arrow in S14). To exclude such noisy voxels, we used a highly conservative approach by thresholding *r*^2^. However, a consequence of this approach is that most of the remaining voxels are located close to the pial surface, where the fMRI signal is strongest, and none or very few are located at the grey matter-white matter boundary. To ensure that all cortical depth layers were represented in the ROIs, and to correct for the fragmented nature of the preliminary ROIs, we first projected the pRF-estimated voxels on the cortical surface, created patches-of-interest, and then projected them back into volume space, where we segmented the ROIs into 6 cortical depth layers (see **Supplementary Fig. 2**). This approach is anatomically justified, as neurons are organized in hypercolumns, which means that neurons located in the deeper layers process input from the same visual field portions as those stacked on top of them (Hubel, 1982). To further validate this approach, we also checked how the preliminary ROIs would look with lowered *r*^2^-cutoffs. As can be seen in the 3 example subjects shown here, more voxels located in the deeper portions of the grey matter were added to the preliminary ROIs (dark red) when the *r*^2^-cutoff was lowered. This confirms the validity of our approach. There is also the reverse case in which, as a consequence of noise, some voxels’ best model falls outside of the visual field of interest, even when it is anatomically evident that these voxels should represent the visual area of interest, as they are surrounded by other voxels that all represent the visual area of interest (see, for example, the voxel indicated by the yellow arrow in S14). Based on anatomical considerations and cross-checks with the retinotopic maps, these voxels were included in the cortical depth layers of the ROIs. For further validation, we additionally re-computed the SVM classification after excluding these voxels. The pattern of results remained the same (data not shown).

**Supplementary Fig. 4.**
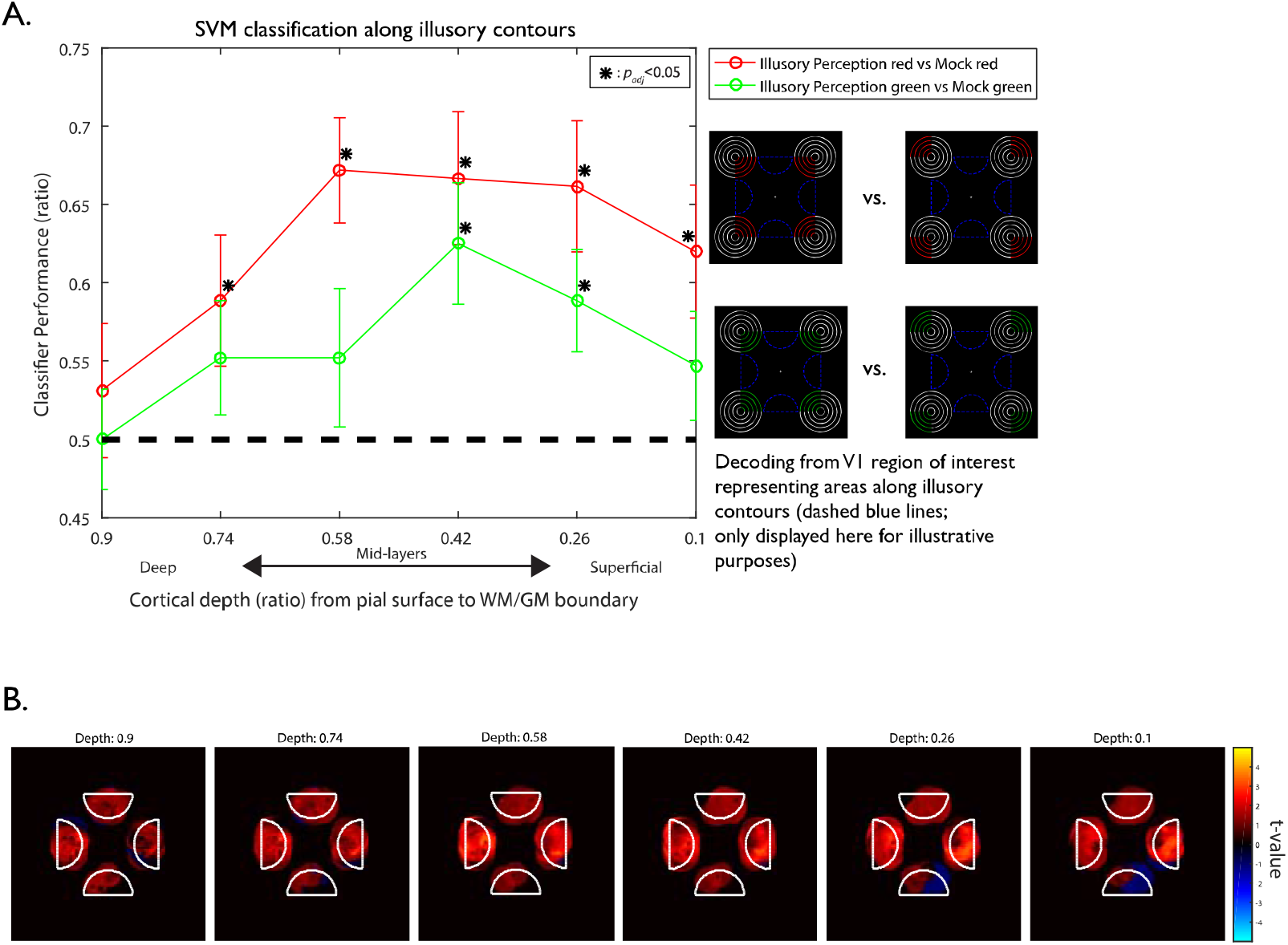
Decoding of illusory perception vs no illusory perception. **A.** We ran additional SVM classification analyses at different cortical depths in the peripheral region of interest (ROI), decoding illusory perception against its mock version, where no illusion should arise. This was done to better compare our classification results with the univariate results of Kok et al.(*9*), who looked at activity differences between illusory and no illusory perception at different V1 depths. As we had illusory and mock stimuli in two colours, we could compute SVM classification analysis twice. The dashed black line designates chance level; asterisks denote significant above-chance decoding (*p*_adj_<.05, FDR-corrected). For both colours, decoding performance increased towards the superficial depths in the peripheral ROI; and significant above-chance decoding was present in both colours at the 2^nd^ and 3^rd^ most superficial depth. This result is in line with our main finding that illusory colour can only be decoded at the second superficial depth. It also matches previous findings that edges between figure and ground selectively activate layer 2/3 in non-human primates(*50*) and mice(*51*). However, it is not in line with Kok’s et al.’s conclusions that illusory perception selectively involves deep layers. **B.** Visual field projections of each voxel’s influence on SVM classification analysis. Instead of displaying voxel influence in cortical space, the visual projection approach translates voxel influence into visual space, giving an idea of which visual field portions (or pixels) contribute most to SVM classification. Shown here are the projections from the SVM classification of the green illusory perception vs. the green mock condition. Deeper depths or shown towards the left, more superficial depths are shown towards the right. We focused on the visual projections of the peripheral ROI voxels, whose pRF centre fell within the peripheral visual field areas of interest (indicated by white outlines). Colour-encoded t-values indicate each visual field pixel’s influence on SVM classification across subjects. The head coil limited visibility in the lower and upper central part of the screen in several participants; for this reason, there was less voxel coverage of these visual field portions, leading to overall weaker t-values here. The visual projections shown here are for visualization purposes; statistical analyses were conducted with the SVM classification analyses shown in A.

**Supplementary Fig. 5.**
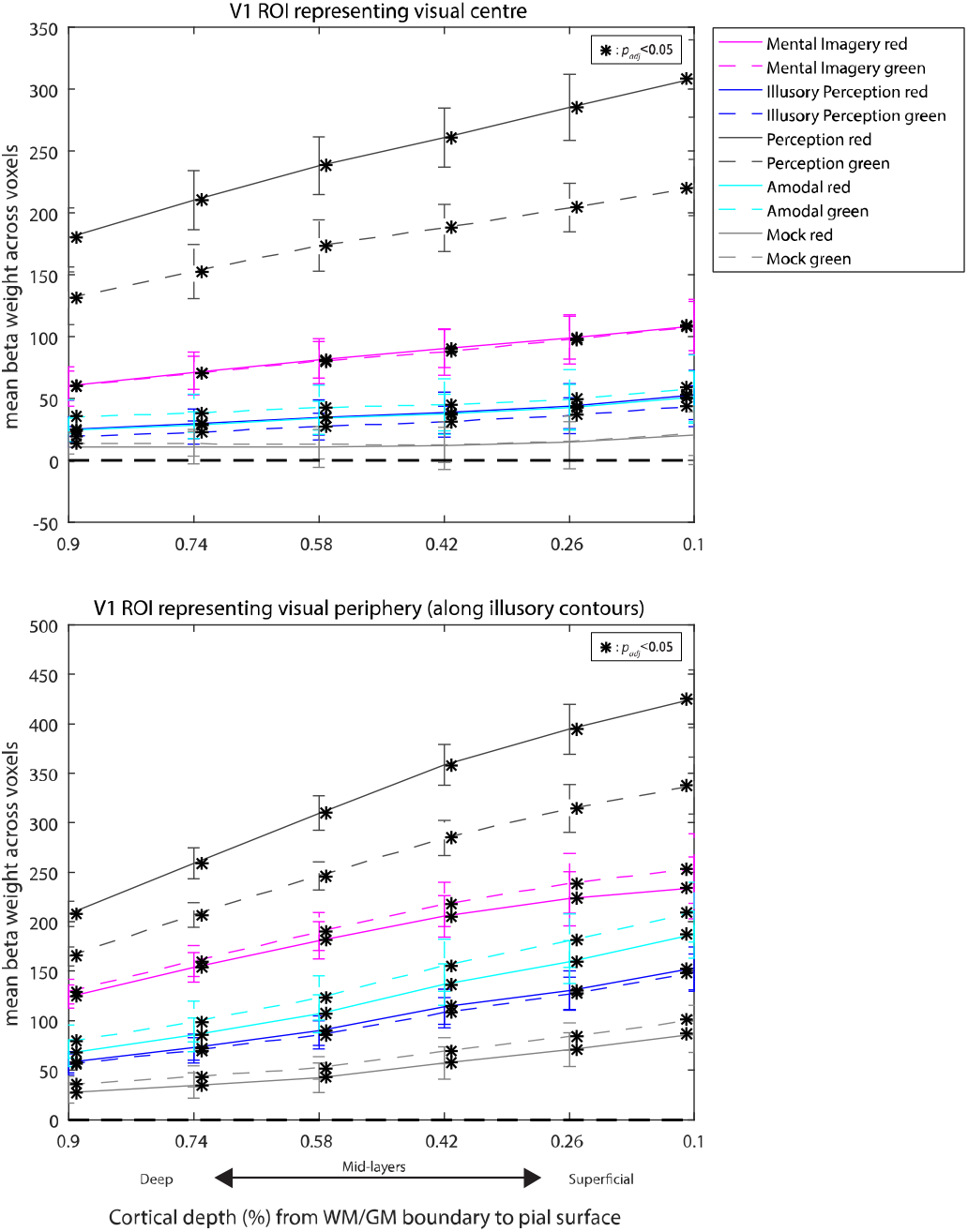
Univariate analysis in the two V1 regions of interest. Univariate analysis across the cortical depth layers of the regions of interest (ROI) that represent the visual centre (upper plot) and the visual periphery (lower plot; also see **Fig. 2C** for a visualization of the visual portions they represent). The cortical depth of 0.9 designates the deepest depth at the grey matter/white matter boundary, and 0.1 designates the most superficial depth at the pial surface. Error bars denote ±SEM. For each depth layer, we computed a GLM analysis (see Methods). The beta weights were then averaged across voxels for each participant, and across participants. The dashed black lines designate a mean beta weight of zero for reference; asterisks denote significant above-chance decoding (*p*_adj_<.05, FDR-corrected). In the visual centre, fMRI BOLD activity (as determined by beta weights) was significantly above zero in all conditions (all *p*_adj_ ≤ .008) except for in the mock condition. In this condition, only the green mock condition was significant, and this was only the case at a cortical depth of 0.9 (*p*_adj_ = .045). In the ROI representing the visual periphery, fMRI activity of all conditions were significantly above zero in all conditions and at all depths (all *p*_adj_ ≤ .001). In a generalized linear mixed effects analysis, the main effects of cortical depth (*χ*^2^(5)=92.87, p<.001), V1 location *χ*^2^(1)=14.84, *p*<.001) and stimulus condition *χ*^2^(9)=67.35, p<.001) were significant, indicating that fMRI activity (1) increases towards superficial depths; (2) was higher overall in the peripheral ROI compared to the foveal ROI; and (3) differed between the different stimulus conditions. In addition, all three 2-way interactions showed significance (all *p*<.01), whereas the 3-way interaction between cortical depth, V1 location and stimulus conditions was not significant *χ*^2^(45)=0, p=1). Also see **Supplementary Fig. 8** and **9** for individual subject plots.

**Supplementary Fig. 6.**
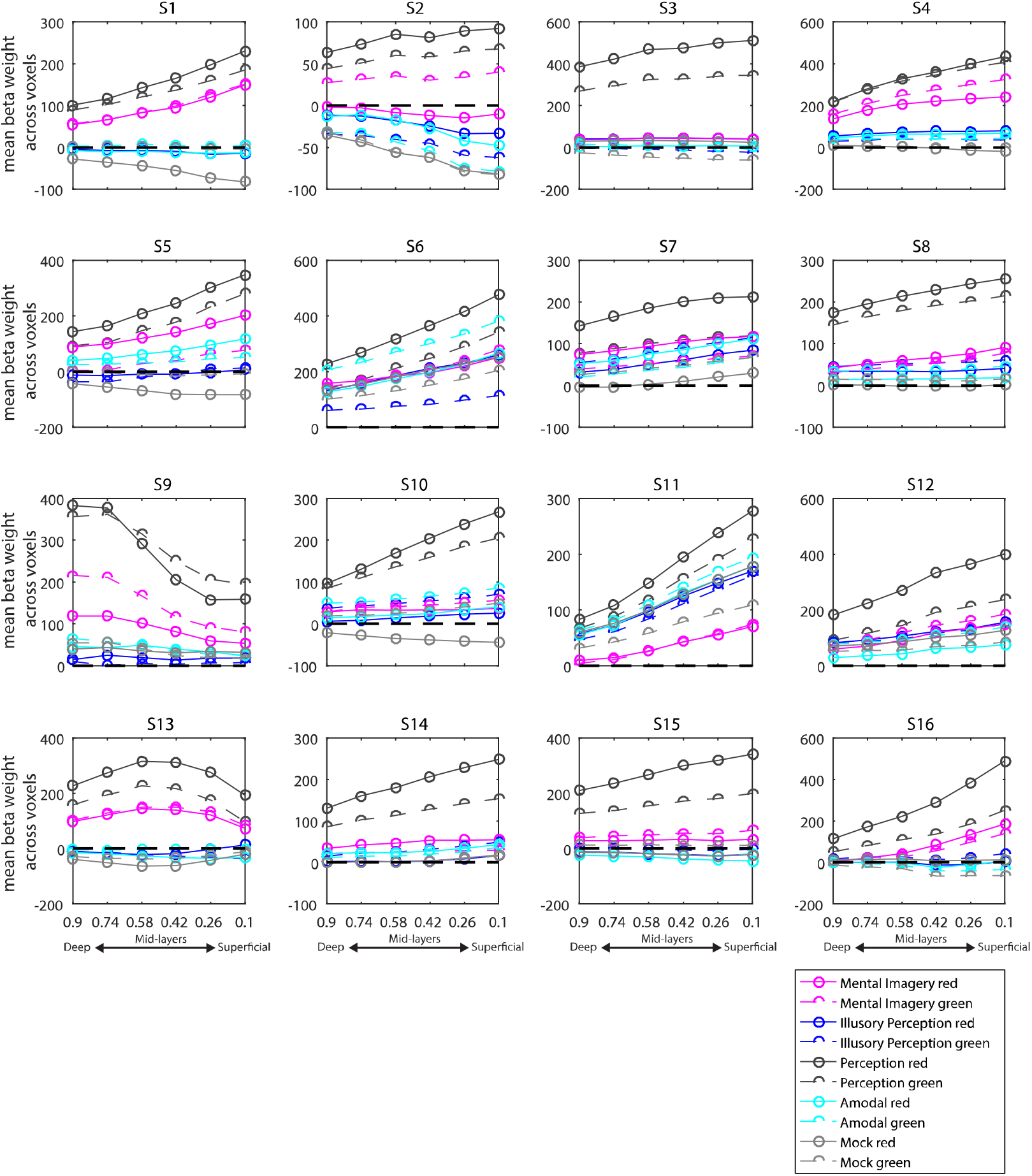
Individual fMRI activity plots in the foveal (central) region of interest. In each plot, deeper depth layers (towards the grey matter/white matter boundary) are shown towards the left, and superficial depth layers towards the right (towards the pial surface). Codes above each plot denote the individual subject. Beta weights for each condition were computed in a GLM analysis and then averaged across voxels. The dashed black lines indicate a mean beta weight of zero for reference.

**Supplementary Fig. 7.**
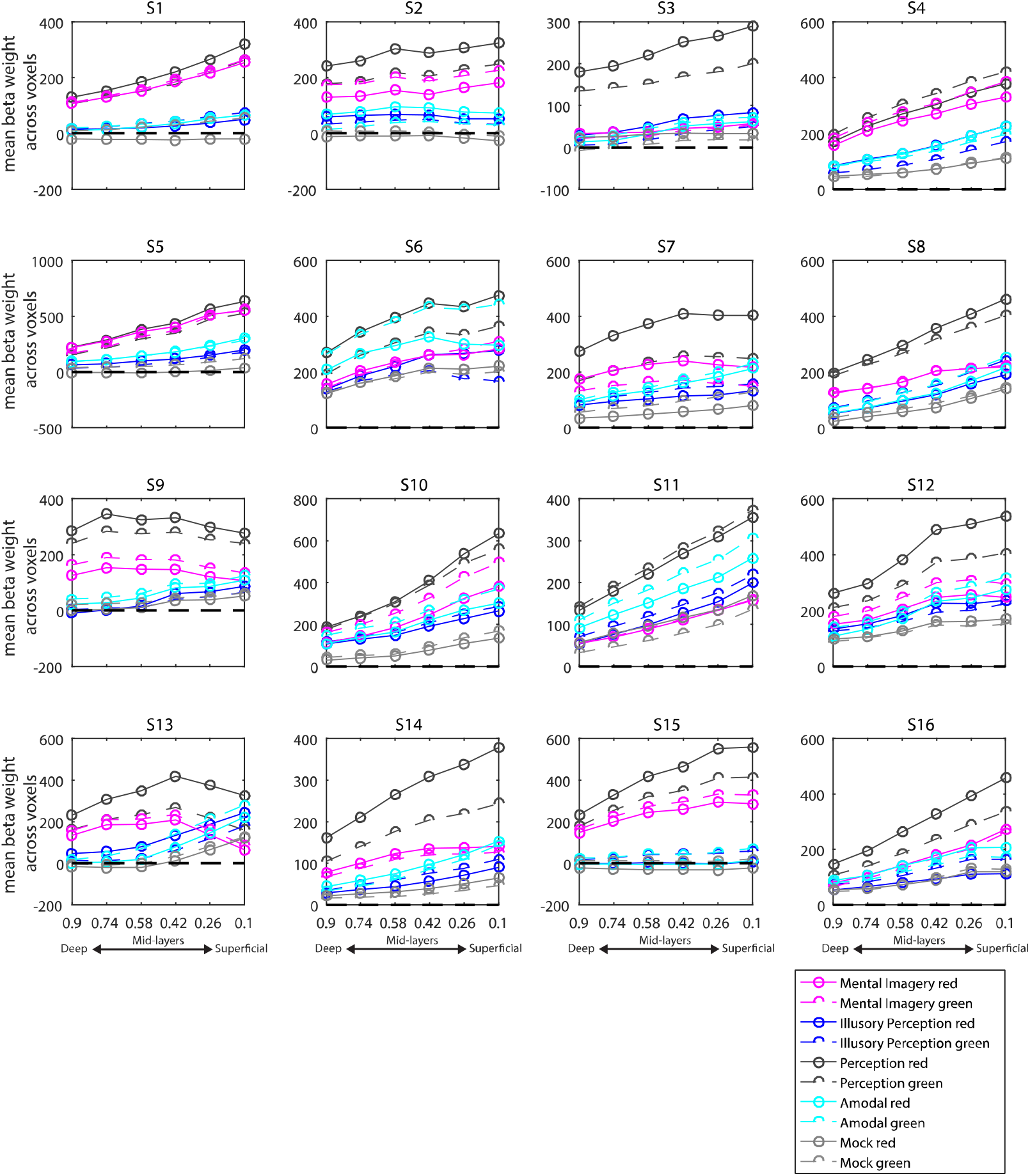
Individual fMRI activity plots in the peripheral region of interest (ROI). In each plot, deep depth layers (towards the grey matter/white matter boundary) are shown on the left, and superficial depth layers on the right (towards the pial surface). Codes above each plot denote the individual subject. Beta weights for each condition were computed in a GLM analysis and then averaged across voxels. The dashed black lines indicate a mean beta weight of zero for reference.

**Supplementary Fig. 8.**
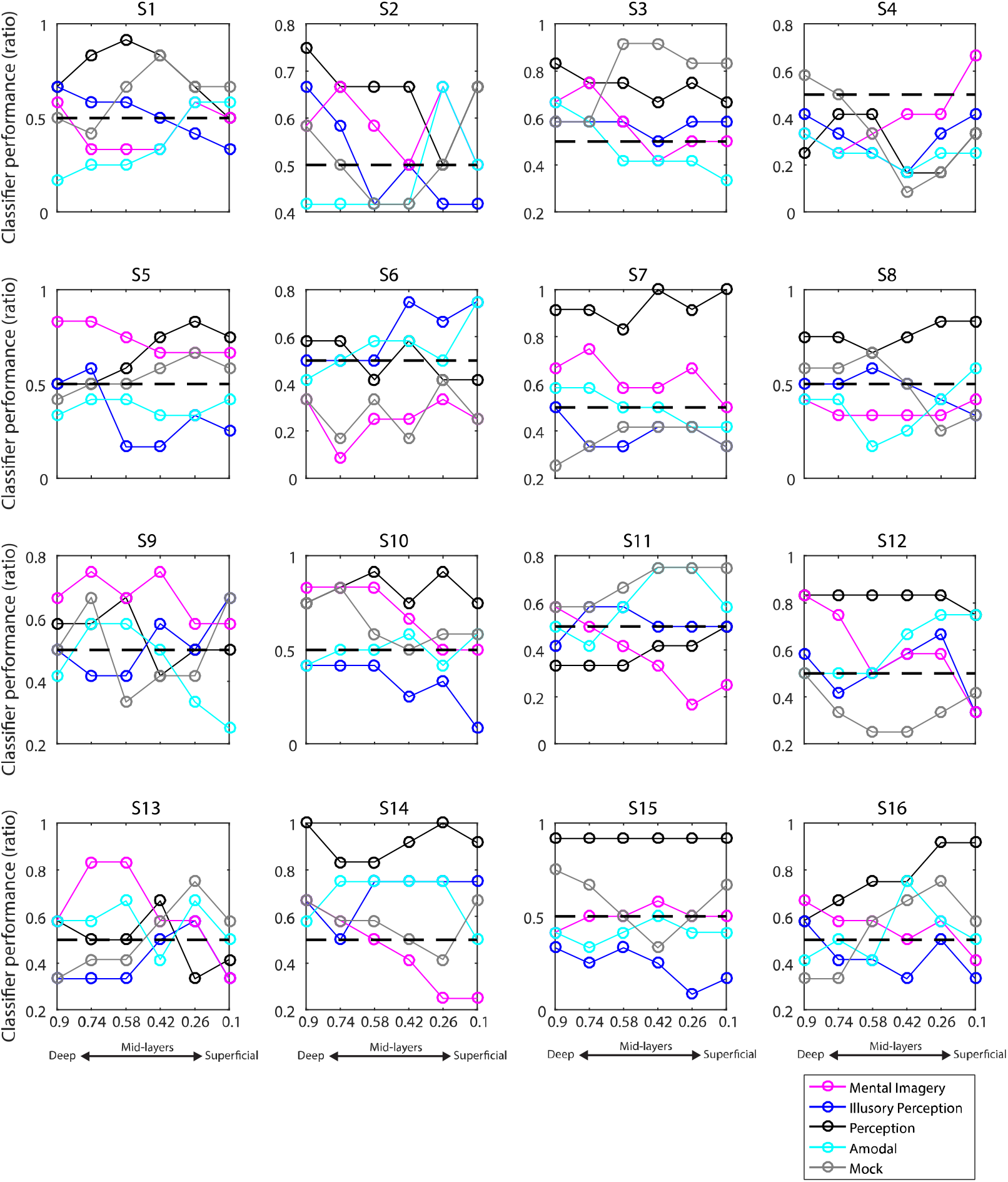
Individual decoding plots in the foveal ROI that represents the central portion of the visual field. Each plot represents the SVM classification data of one participant. In each plot, deeper depth layers (towards the grey matter/white matter boundary) are shown towards the left, and superficial depth layers towards the right (towards the pial surface). Chance level is indicated by the dashed black lines. Codes above each plot denote the subjects.

**Supplementary Fig. 9.**
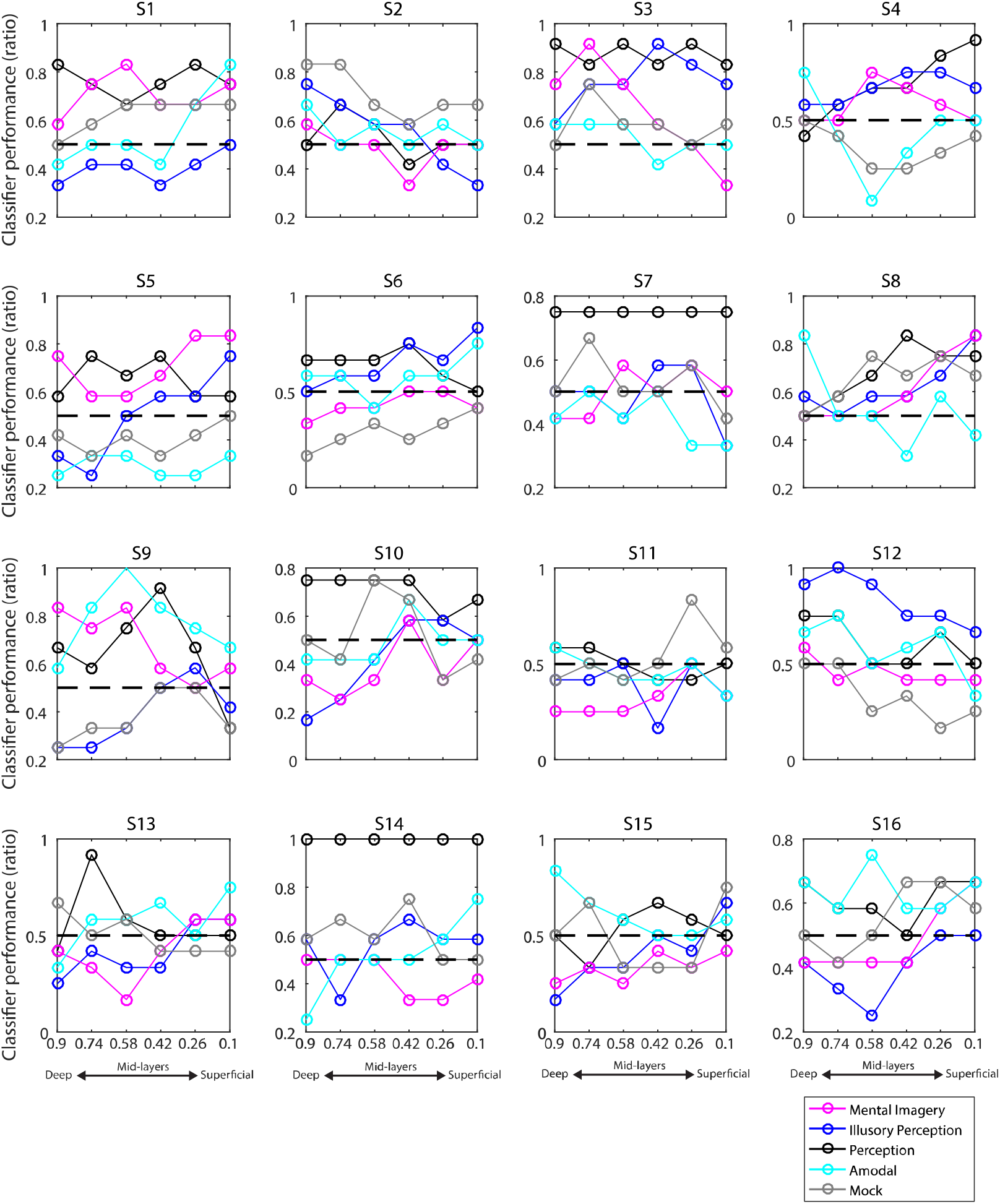
Individual decoding plots in the peripheral ROI that represents the peripheral portions of the visual field. Each plot represents the SVM classification data of one participant. In each plot, deeper depth layers (towards the grey matter/white matter boundary) are shown towards the left, and superficial depth layers towards the right (towards the pial surface). Chance level is indicated by the dashed black lines. Codes above each plot denote the subjects.

